# The SspB adaptor drives structural changes in the AAA^+^ ClpXP protease during ssrA-tagged substrate delivery

**DOI:** 10.1101/2022.11.06.515074

**Authors:** Alireza Ghanbarpour, Xue Fei, Tania A. Baker, Joseph H. Davis, Robert T. Sauer

**Author notes:** Co-first authors (Alireza Ghanbarpour, Xue Fei). Equal contributions (Joseph H. Davis, Robert T. Sauer). **Classification:** Biological Sciences; Biophysics and Computational Biology.

## Abstract

Energy-dependent protein degradation by the AAA^+^ ClpXP protease helps maintain protein homeostasis in organisms ranging from simple bacteria to humans. In *E. coli* and many other proteobacteria, the SspB adaptor assists ClpXP in degrading ssrA-tagged polypeptides produced as a consequence of tmRNA-mediated ribosome rescue. By tethering these incomplete ssrA-tagged proteins to ClpXP, SspB facilitates their efficient degradation at low substrate concentrations. How this process occurs structurally is unknown. Here, we present a cryo-EM structure of the SspB adaptor bound to a GFP-ssrA substrate and to ClpXP. This structure provides evidence for simultaneous contacts of SspB and ClpX with the ssrA tag within the tethering complex, allowing direct substrate handoff concomitant with the initiation of substrate translocation. Furthermore, our structures reveal that binding of the substrate•adaptor complex induces unexpected conformational changes within the spiral structure of the AAA^+^ ClpX hexamer and its interaction with the ClpP tetradecamer.

**SIGNIFICANCE:** Intercellular proteases, including ClpXP, degrade damaged or unneeded proteins. Peptide tags allow specific protein substrates to be recognized by the ClpX unfoldase/translocase component of ClpXP and by an adaptor, SspB, which tethers itself to ClpX and enhances ClpXP degradation of the tagged protein. Our cryo-EM structure of ClpXP bound to SspB and a tagged substrate shows that SspB and ClpX simultaneously contact the degradation tag and reveal changes in the structure of ClpX and its interaction with ClpP. These structural changes appear to be a prelude to an initial ClpX translocation step that pulls the substrate away from SspB and initiates degradation by allowing substrate unfolding and further translocation of the unfolded substrate into the proteolytic chamber of ClpP.

## INTRODUCTION

Proteolytic adaptors alter the repertoire of substrates degraded by individual proteases (Mahmoud and Chien, 2018). For example, in *Escherichia coli*, the SspB adaptor changes the proteolytic fate of ssrA-tagged proteins, which are produced when protein biosynthesis on a ribosome stalls and the tmRNA system completes translation by attaching an ssrA tag to the C-terminus of the partial protein (Keiler *et al*., 1996; Levchenko *et al*., 2000). Translation can stall at many different codons in any protein-coding segment of the *E. coli* genome, and thus ssrA-tagged proteins are highly diverse in terms of sequence, size, and structure (Moore and Sauer, 2007). Importantly, the ssrA tag serves as a degron for the cytoplasmic AAA^+^ ClpXP and ClpAP proteases (Gottesman *et al*., 1998). Thus, any ssrA-tagged protein in the *E. coli* cytoplasm can potentially be degraded, ridding the cell of truncated, useless, and potentially dangerous protein fragments and permitting the amino acids in these aberrant molecules to be recycled.

Each subunit of the SspB homodimer consists of a native domain, which binds to the first seven residues of the ssrA tag (<u>AANDENY</u>ALAA), and a disordered C-terminal region, ending with a short segment that binds to the N-domain of ClpX (Bolon *et al*., 2004a; Chien *et al*., 2007; Levchenko *et al*., 2003; Song and Eck, 2003; Wah *et al*., 2003). These SspB-mediated interactions tether ssrA-tagged proteins to ClpXP and stimulate their degradation by decreasing *K*_M_ and increasing V_max_ (Wah *et al*., 2003; Chien *et al*., 2007). Concurrently, SspB redirects degradation by blocking recognition and proteolysis of ssrA-tagged proteins by the ClpAP protease (Flynn *et al*., 2001).

ClpXP consists of one or two hexamers of ClpX and the double-ring ClpP tetradecamer (Baker and Sauer, 2012). The ClpX ring is assembled by packing of six AAA^+^ modules of the hexamer in a shallow spiral around an axial channel that serves as a conduit to the degradation chamber of ClpP (Fei *et al*., 2020a; Fei *et al*., 2020b; Ripstein *et al*., 2020). ClpX^ΔN^•ClpP, which lacks the N-terminal domain, which binds to SspB, is active in degradation of some protein substrates (Singh *et al*., 2001; Martin *et al*., 2005), including ssrA-tagged proteins, and has been used for most prior cryo-EM studies (Fei *et al*., 2020a; Fei *et al*., 2020b; Ripstein *et al*., 2020). Early biochemical experiments established that positively charged “RKH” loops that reside at the top of the ClpXP axial channel aid in initial ssrA tag recruitment (Farrell *et al*., 2007). It is hypothesized that following recruitment, the C-terminal ALAA residues of the ssrA tag bind in the upper portion of a closed ClpX channel, as shown in a“recognition-complex” structure (Fei *et al*., 2020a). Subsequent channel opening would then be needed to allow tag translocation and eventual substrate unfolding. Following this multistep ssrA tag recognition, ClpX or ClpX^ΔN^ use translocation steps powered by ATP hydrolysis to unfold any native structure present in the substrate and to spool the unfolded polypeptide through the axial channel and into ClpP for degradation (Baker and Sauer, 2012; Sauer *et al*., 2022).

The structural manner in which SspB interacts with ClpXP and stimulates substrate degradation is poorly understood. For example, when ClpX-tethered SspB releases the ssrA-tagged substrate relative to tag recruitment, tag recognition, and pore opening of the ClpX channel is unclear. Moreover, the molecular mechanisms by which adaptors stimulate degradation by AAA^+^ proteases is understudied, with only the minimal requirements having been established (Davis *et al*., 2009; 2011).

Here, we present a cryo-EM structure of a ClpXP•GFP-ssrA•SspB (hereafter protease•substrate•adaptor) complex that demonstrates that SspB directly positions the C-terminal residues of the ssrA tag in the upper part of a closed axial channel of ClpX, almost exactly as in an adaptor-free recognition complex (Fei *et al*., 2020a). These simultaneous contacts between SspB, ClpX, and the ssrA tag allow direct hand-off of the substrate from the adaptor to the protease. Surprisingly, we also find that binding of the substrate•adaptor causes a substantial alteration in the spiral arrangement of ClpX subunits and how they interact with ClpP. We discuss the implications of this latter finding for the mechanism by which ClpXP and other AAA^+^ proteases engage protein substrates as a prelude to ATP-fueled unfolding and eventual degradation.

## RESULTS AND DISCUSSION

### Structure of the protease•substrate•adaptor complex

We combined *E. coli* ClpX, ClpP, SspB, and *Aequorea victoria* green-fluorescent protein with an appended *E. coli* ssrA tag (GFP-ssrA) in the presence of ATP_ϒ_S and determined a cryo-EM structure of the complex (Supplementary Table 1, Supplementary Figures 1–3). ATP_ϒ_S supports assembly of ClpXP and binding to SspB and GFP-ssrA but is hydrolyzed too slowly to support GFP-ssrA unfolding and degradation (Wah *et al*., 2002; Martin *et al*., 2008b).

In our cryo-EM structure, we observed density for the large and small AAA^+^ domains of ClpX, for the ssrA tag, for both heptameric rings of ClpP, and for the two native domains of SspB, although density for the one proximal to ClpX was better resolved (Figure 1). By contrast, the ClpX N-domains, the C-terminal residues of SspB, and the native barrel of GFP were not visible in the density map, presumably as a consequence of these structures occupying multiple orientations given that the ClpX N-domains are flexibly joined to the AAA^+^ modules and a short unstructured region separates native GFP from the ssrA tag. ATP_ϒ_S or ADP was bound at the interfaces between the large and small AAA^+^ domains of all six ClpX subunits.

**Figure 1.**
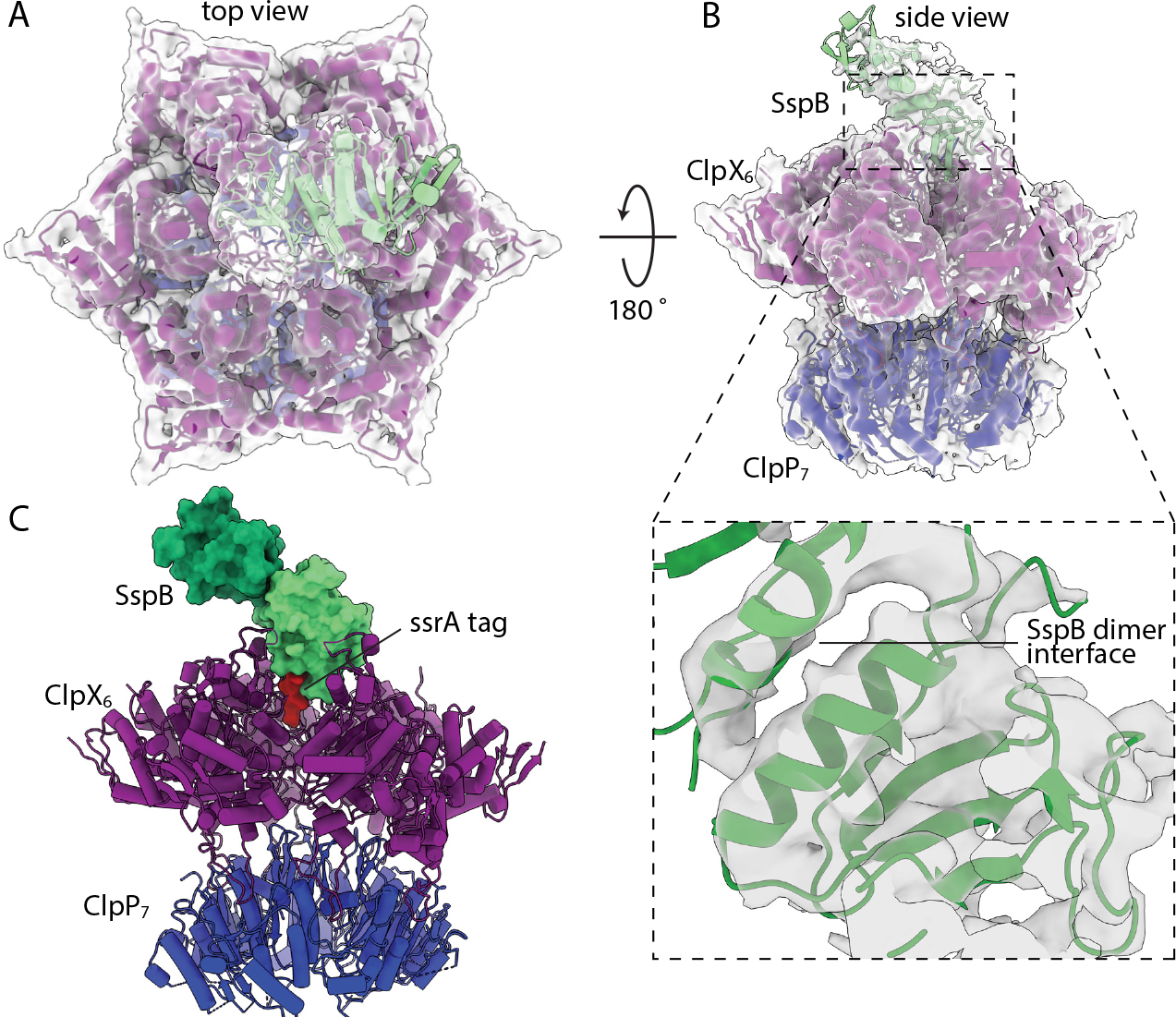
Protease•substrate•adaptor cryo-EM desnity map and atomic model. Overlay of density map (gray) and model (cartoon representation) of the protease•substrate•adaptor structure in top (**A**) and side (**B**) views. The inset in panel B depicts the map and model for the proximal SspB subunit and dimer interface. (**C**) Atomic model of the complex. The native domains of the proximal and distal SspB subunits are shown in surface representation and colored light and dark green, respectively. The ssrA tag is shown in surface representation and colored red. ClpX is shown in cartoon representation and colored purple. The heptameric ring of ClpP nearest ClpX is shown in cartoon representation and colored blue.

### Protease and adaptor simultaneously contact substrate during delivery

In our protease•substrate•adaptor structure, the SspB subunit proximal to ClpX positioned the ssrA tag to make bridging interactions with the top portion of a closed axial channel of ClpX (Figure 2). This bridging geometry in the protease•substrate•adaptor delivery complex would allow subsequent opening of the axial channel and complete transfer of the substrate to the protease. For example, an ATP-fueled ClpX translocation step could open the channel by pulling the ssrA tag deeper into the pore. Such a motion could simultaneously release the substrate from the SspB adaptor, which because of its bulk could not enter the channel. Subsequent translocation steps would then result in substrate unfolding and degradation.

**Figure 2.**
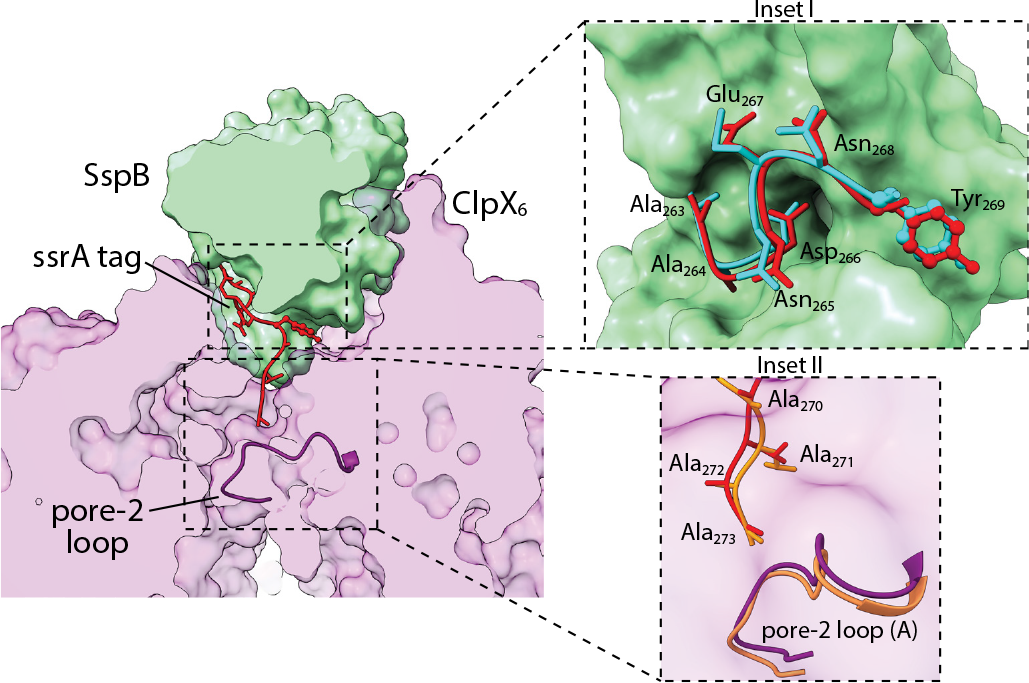
SspB • ssrA and ClpX • ssrA contacts. Substrate delivery complex clipped in plane to highlight contacts between the proximal subunit of SspB (green surface representation), the ssrA tag (red stick or ball-and-stick representation), and ClpX (light purple; surface representation). The pore-2 loop (dark purple backbone representation) of chain A of ClpX blocks the axial channel and contacts the C-terminus of the ssrA tag. **Inset I** highlights SspB contacts with the ssrA tag. The conformation of an ssrA tag from a crystal structure (pdb 1OX9) bound to SspB (cyan stick or ball-and-stick representation) is also shown. **Inset II** depicts contacts between ClpX and the C-terminal residues of the ssrA tag overlaid with models of the same tag residues and pore-2 loop (orange) from the recognition complex (pdb 6WRF).

The seven N-terminal residues of the ssrA tag in our cryo-EM structure had essentially the same conformation and contacted SspB in the same manner observed previously in an ssrA•SspB crystal structure (Song and Eck, 2003) (Figure 2). Moreover, the four C-terminal residues of the tag were positioned similarly in our current structure and in the recognition-complex structure (Fei *et al*., 2020a), further supporting a bridging followed by a handoff mechanism. In both the new structure and recognition complex, the pore-2 loop of ClpX subunit A closed the axial channel and contacted the C-terminus of the ssrA tag (Figure 2). Mutation of the pore-2 loop weakens ClpXP affinity for a crosslinked ssrA-SspB complex more than 60-fold (Martin *et al*., 2008a), and thus the contacts between the pore-2 loop and the ssrA tag in our structure make substantial contributions to the thermodynamic stability of the protease•substrate•adaptor complex. SspB and ClpX also contacted opposite faces of the aromatic ring of the single tyrosine in the ssrA tag in the structure (Figure 3), emphasizing the very tight packing in this region. This close packing may result in minor steric or electrostatic clashes, and could explain why inserting several residues in this region of the tag strengthens ssrA•SspB binding to ClpX (Hersch *et al*., 2004).

**Figure 3.**
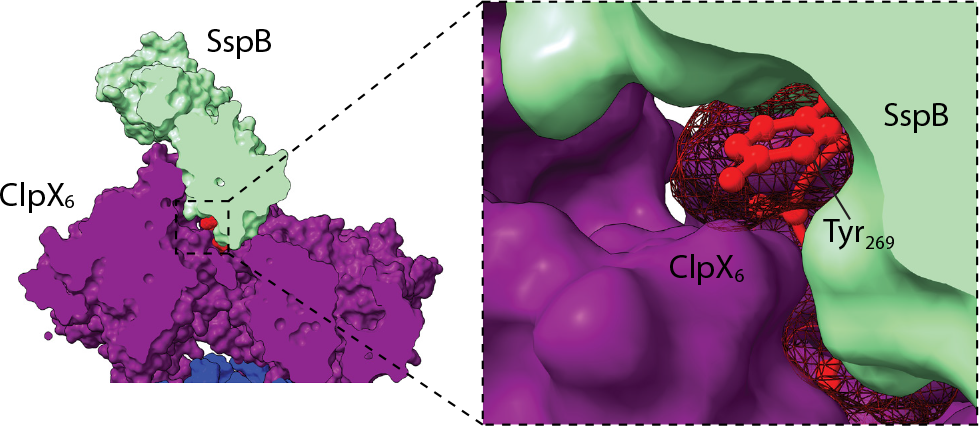
The SspB-ssrA-ClpX interface. SspB and ClpX contact opposite sides of the aromatic ring of the tyrosine in the ssrA tag highlighted in the mesh surface representation.

### RKH loops contact SspB but provide little stabilization

ClpX has the axial pore-1 and pore-2 loops common to all proteolytic AAA^+^ enzymes but also contains family specific RKH loops, which surround the entrance to the translocation channel and play roles in substrate and adaptor binding and specificity (Martin *et al*., 2008a; Baker and Sauer, 2012; Fei *et al*., 2020a; Ripstein *et al*., 2020). In our structure, multiple ClpX RKH loops contacted the proximal SspB subunit (Figure 4), as anticipated by crosslinking results (Martin *et al*., 2008a).

Interestingly, however, our covariation analysis using RaptorX (Ma *et al*., 2015) or GREMLIN (Ovchinnikov *et al*., 2014) showed no significant mutual-sequence information between ClpX and SspB. Moreover, only the dimer interface and ssrA-binding groove of SspB showed high levels of sequence conservation, whereas most residues contacted by ClpX in our structure were poorly conserved (Figure 4). These findings and the very low affinity of tail-less SspB dimers for ClpX (Bolon *et al*., 2004b) suggest that the interactions between the RKH loops and SspB are roughly neutral with respect to free energy, with favorable interactions cancelled by unfavorable interactions or entropic costs. By this model, SspB binding is stabilized mainly by its tethering to the N-domains of ClpX and by the bridging of the ssrA-tag between SspB and ClpX. Relatively weak binding of substrate-free SspB to ClpX could be biologically important in avoiding competition between substrate-free SspB adaptors and the degradation of substrates without ssrA tags that need unrestricted access to the ClpX channel.

**Figure 4.**
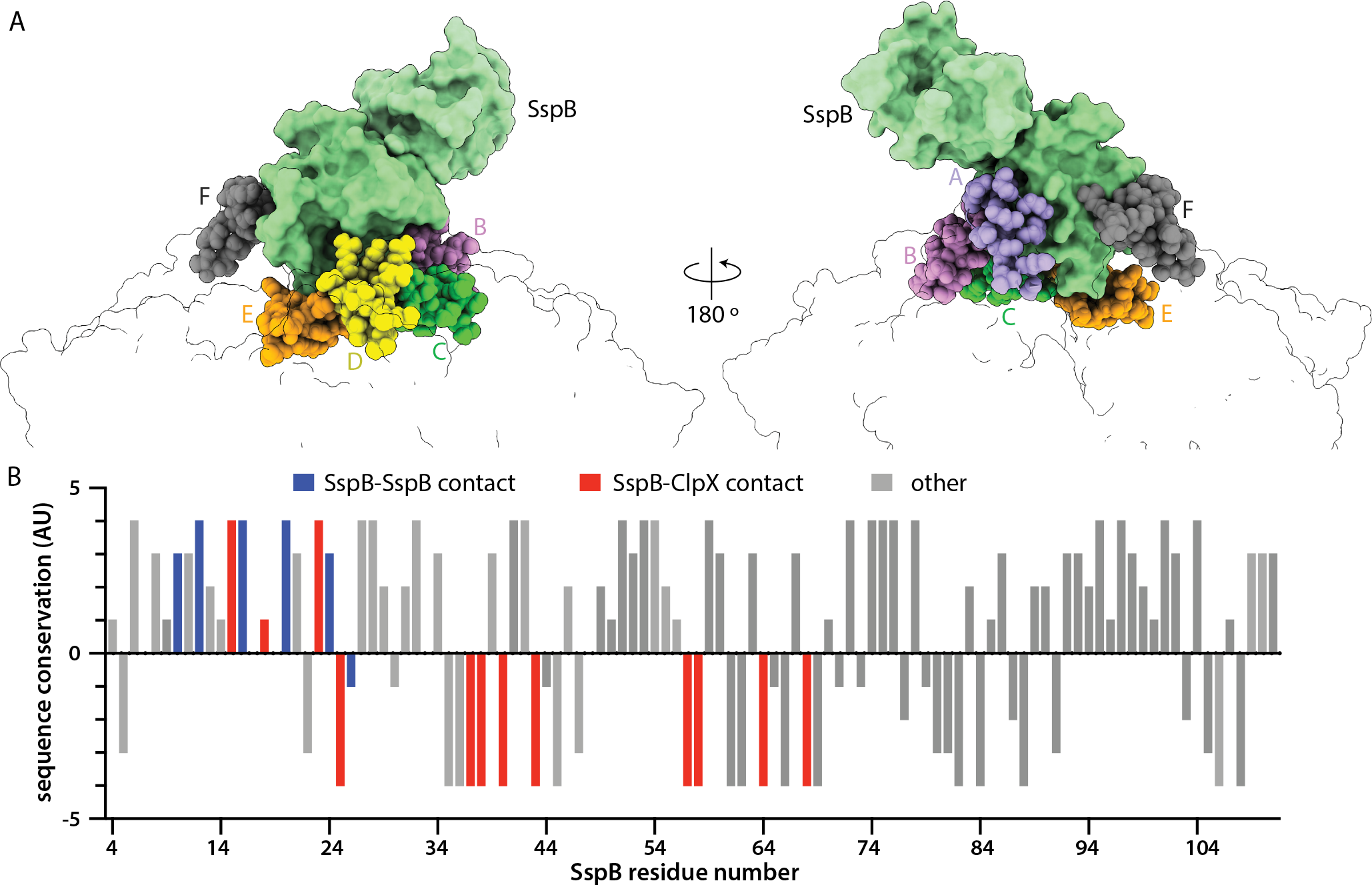
SspB contacts with ClpX RKH loops are poorly conserved. (**A**) The folded domain of SspB proximal to ClpX (green surface representation) is contacted by six RKH loops (sphere representation colored by subunit identity). (**B**) Conservation score of SspB residues calculated by Consurf (Ashkenazy et al., 2016) plotted against sequence position. Positive values indicate higher sequence conservation. Residues classified and bars colored based on SspB residue contacts. Gray bars (other) note residues lacking ClpX or SspB inter-dimer contacts. For this analysis, SspB residues within 5 Å from interacting molecules were considered as contacts.

### Conformational changes in ClpXP are induced by substrate•SspB binding

In our protease•substrate•adaptor complex, the large AAA^+^ domains of ClpX subunits A, B, C, D, and E formed a shallow spiral, as observed in previous ClpXP structures (Fei *et al*., 2020a; Fei *et al*., 2020b; Ripstein *et al*., 2020). Notably, however, the large AAA^+^ domain of subunit F moved out of the spiral and up towards SspB, as a consequence of a rotation relative to its small AAA^+^ domain (Figure 5). This movement of subunit F was accompanied by smaller movements of subunit E and a minor shift in subunit A. Supplemental movies 1 and 2 show morphs between the protease•substrate•adaptor complex and recognition complex as viewed from subunits A or D.

**Figure 5.**
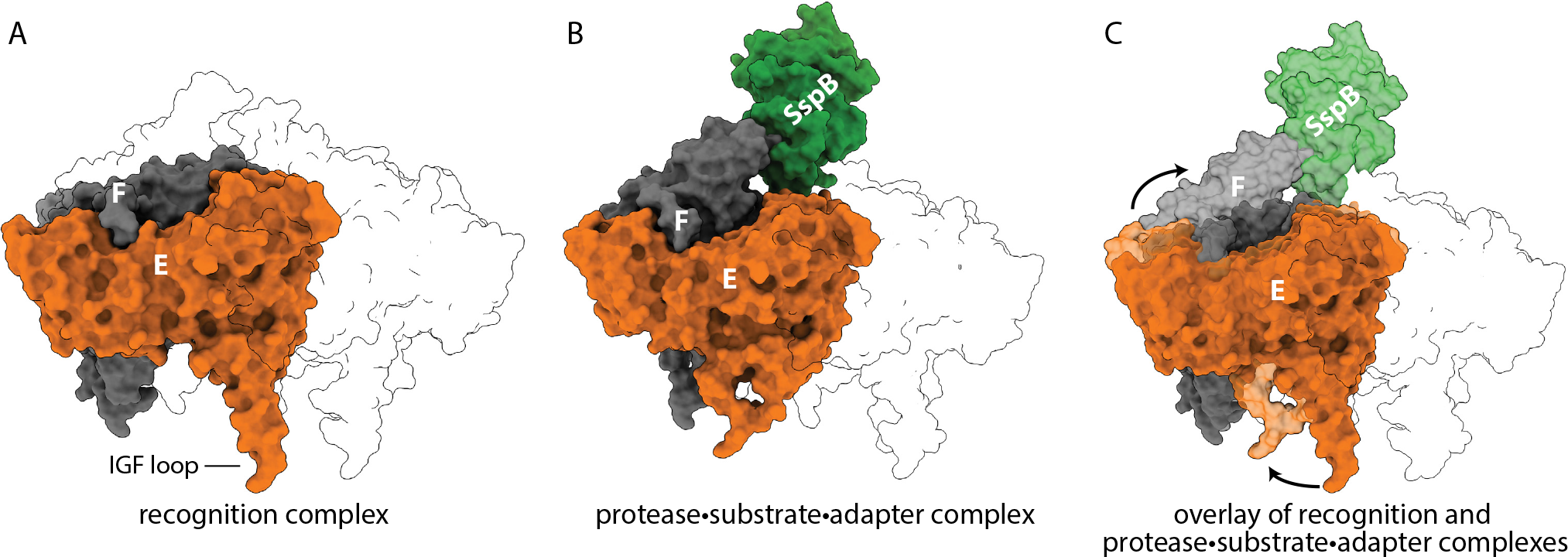
Structural changes in ClpX caused by the binding of GFP-ssrA•SspB. (**A**) ClpX subunits F (gray) and E (orange) are shown in the recognition complex (pdb 6WRF) with other subunits in transparent outline. (**B**) Positions of subunits F and E and SspB (green) in the protease•substrate•adaptor structure. (**C**) Overlay of the recognition complex in darker shades with the protease•substrate•adaptor complex in lighter shades highlights the upward movement of subunit F and leftward movement of subunit E.

These movements of ClpX subunits in the protease•substrate•adaptor complex resulted in changes in the contacts between ClpX and ClpP. The six IGF loops of a ClpX hexamer fit into six of the seven docking clefts in a heptameric ring of ClpP, leaving one cleft empty. In prior structures of *E. coli* ClpXP (Fei *et al*., 2020a; Fei *et al*., 2020b), this empty cleft was located between the IGF loops of ClpX subunits E and F. In our protease•substrate•adaptor structure, by contrast, the empty cleft was located between the IGF loops of subunits D and E (Figure 6).

**Figure 6.**
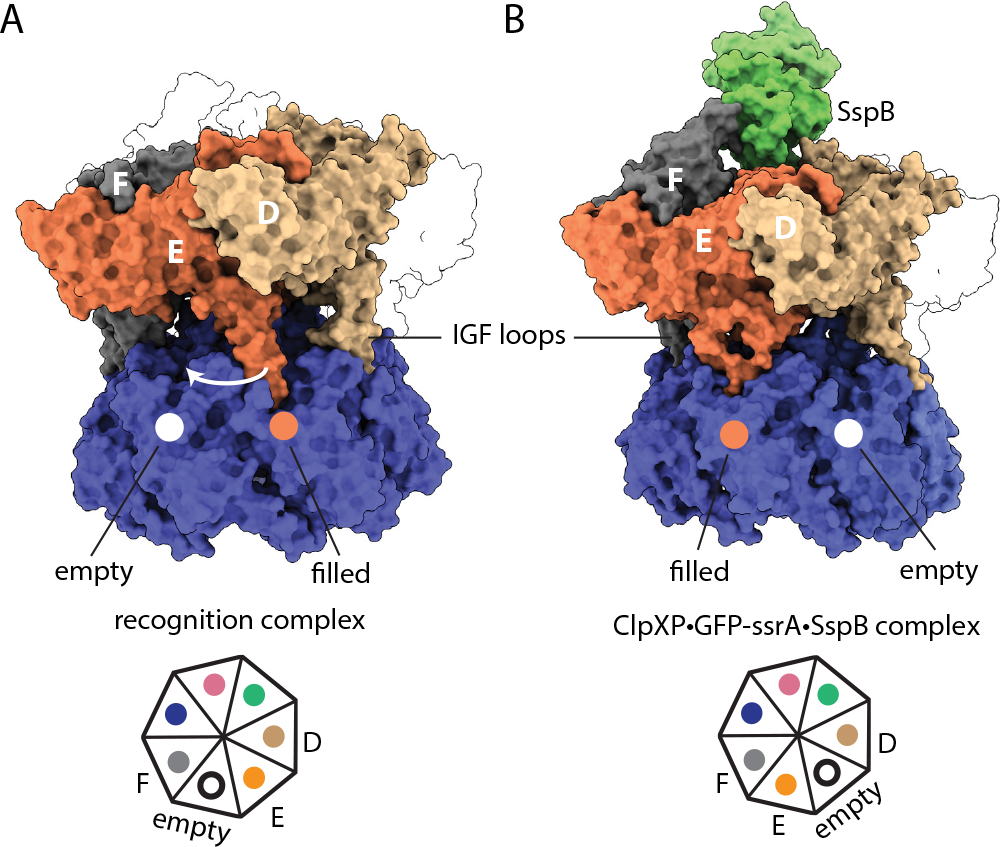
Substrate•adaptor mediated movement of IGF loops. (**A**) The empty ClpP cleft is located between the IGF loops of sub-units E and F in the recognition complex (pdb 6WRF). (**B**) In the protease•substrate•adaptor complex, the empty cleft was located between ClpX subunits E and D.

Does the ClpX conformation observed in our protease•substrate•adaptor structure have a specialized function? Biophysical experiments indicate that ClpXP adopts a kinetic state distinct from a translocation complex state when it tries to unfold a substrate (Saunders *et al*., 2020). Thus, the novel ClpX conformation in our protease•substrate•adaptor complex might correspond to a state capable of applying an efficient unfolding force. Such a conformation could help ClpX to resist the equal-and-opposite force imposed by a distorted substrate during a denaturation attempt and thus improve the chances of successful unfolding. If this surmise is correct, then similar conformations of the ClpX hexamer should be observed in SspB-free but substrate-engaged structures, where ‘substrate-engaged’ is defined as a state in which one additional translocation step would result in substrate denaturation.

### Summary

Our protease•substrate•adaptor structure provides the first glimpse of a critical step in SspB delivery of ssrA-tagged substrates for ClpXP degradation and reveals a new conformation of the ClpX hexamer. It also provides an explanation for the ability of SspB to lower *K*_M_ for ClpXP degradation of ssrA-tagged substrates, namely that ClpX•ssrA contacts are augmented by interactions between ssrA•SspB and SspB•ClpX to increase affinity.

## ACKNOWLEDGMENTS

Supported by NIH grants R01-GM144542, R35-GM141517, and NSF-CAREER grant 2046778. Samples were prepared at the Automated Cryogenic Electron Microscopy Facility in MIT.nano and screened on a Talos Arctica microscope, which was a gift from the Arnold and Mabel Beckman Foundation. We thank Ed Brignole and Sora Kim for advice and assistance.

## CONTRIBUTIONS

Biochemical experiments (AG; XF); cryo-EM and model building (AG; XF; JHD; RTS); supervision (TAB; JHD; RTS); manuscript preparation (AG; JHD; RTS).

## MATERIALS AND METHODS

### Protein expression purification

*E. coli* ClpP, *E. coli* ClpX (containing a neutral K408E mutation), and *E. coli* SspB were expressed and purified as reported (Levchenko *et al*., 2005). *A. victoria* GFP-ssrA was expressed and purified as described (Kim *et al*., 2000).

### Cryo-EM sample preparation and analysis

For cryo-EM sample preparation, ClpX_6_ (5.4 μM), ClpP_14_ (1.8 μM), GFP-ssrA (10 μM), and SspB_2_ (10 μM) were incubated with ATP_ϒ_S (5 mM) in 20 mM HEPES (pH 7.5), 20 mM KCl, 5 mM MgCl_2_, 10% glycerol, and 0.032% Nonidet P40. The sample was plunge frozen on 300-mesh Quantifoil copper grids, which had been glow-discharged for 60 s in an easiGlow glow discharger (Pelco) at 15 mA and blotted using a Vitrobot Mk IV instrument (Thermo Fisher Scientific) for 3 s with a blot force of 10 (25 °C; 100% relative humidity). For the ClpXP•GFP-ssrA•SspB structure, 9,511 movies were collected at a magnification of 105,000 X and detected in super resolution mode on a Gatan K3 detector for an effective pixel size of 0.435 Å (super resolution mode) with EPU (Thermo Fisher Scientific) on a Titan Krios G3i (Thermo Fisher Scientific), operating at an acceleration voltage of 300 kV. Movies were collected as 30 frames with a total exposure on specimen of 75.98 e-/Å^2^. Defocus ranged from −1.0 to −2.5 μm.

Frames in each movie were binned (2-fold), aligned, gain-corrected, and dose-weighted using Motioncor2 (Zheng *et al*., 2017) to generate micrographs. The contrast transfer function (CTF) was estimated using CTFFIND4 (Rohou and Grigorieff, 2015). RELION 3.1 (Zivanov *et al*., 2018) was used for 2D/3D classification and refinement (see Supplementary Figures 1,2). After several rounds of 2D classification, particles were re-extracted and combined using the join star tool in RELION, and the resulting 391,668 particles were selected for 3D reconstruction.

First, particles were subjected to a 3-class 3D classification with pose estimation using a 40-Å low-pass filtered ClpP map (EMDB: EMD-20434) as the initial model. No masks were used at this stage. The best resolved class contained 236,728 particles. These particles underwent auto-refinement without symmetry (C1) and post processing yielded a ClpXP map at 3.8 Å resolution. Three rounds of CTF-refinement and particle polishing followed by an additional round of 3D auto-refinement improved the overall resolution of this map to 3.2 Å.

Next, a mask encompassing SspB, ClpX, and the cis-ring of ClpP was generated and particle subtraction was applied to the particle stacks to focus on this region of the map. Relion reconstruct was applied to this subtracted particle stack to generate an initial model for a 3-class 3D classification with pose estimation. A final round of 3D auto-refinement on the best resolved 3D class (236,728 particles) and post processing produced map focused on SspB•ClpX•ClpP_cis-ring_ at a resolution of 3.7 Å. This map and associated half-maps were then rescaled using a calibrated pixel size (0.416 Å) that was determined by analyzing the real space correlation between publicly available apo-ferritin density maps and maps of apo-ferritin determined using this microscope under identical imaging condition. The final rescaled maps were used to estimate local and global resolution, as well as directional resolution (Tan *et al*., 2017) (Supplementary Figure 3).

To build atomic models, ClpX, ClpP, and SspB structures (pdb codes 6PP8, 6PPE, and 1OX9) were docked into EM maps using ChimeraX-1.3 (Pettersen *et al*., 2021). ClpX domains were rigid body refined using Coot (Emsley and Cowtan, 2004), and real-space refinement was performed using Phenix 1.14 (Adams *et al*., 2010). The ssrA degron was modeled and refined using Coot and Phenix. All model building relied on the rescaled, but unsharpened maps noted above.

## DATA SHARING PLAN

The structure has been deposited in the Protein Data Base (pdb entry 8ET3) and the cryo-EM density map has been deposited in the Electron Microscopy Data Base (emdb entry 28585). The electron micrographs are being deposited in the Electron Microscopy Public Image Archive.

---

**Supplementary Table 1.** Cryo-EM data collection, processing, model building, and validation statistics.

| Sample and data deposition information |  | Model composition |  |
| --- | --- | --- | --- |
| Nucleotide added | ATPyS | Non-hydrogen atoms | 53,287 |
| PDB ID | 8ET3 | Protein residues | 3,639 |
| EMDB ID | 28585 | Ligands | 4 ATPyS, 2 ADP |
| EMPIAR ID | to be deposited |  |  |
| Data collection |  | Model refinement |  |
| Microscope | Titan Krios G3i | Refinement package | Phenix |
| Camera / mode | Gatan K3 / counting | Map-to-model cross correlation | 0.78 |
| Magnification (nominal) | 105,000 X | RMS deviation bond lengths (Å) | 0.002 |
| Accelerating voltage (kV) | 300 | RMS deviation bond angles (°) | 0.633 |
| Total electron dose (e<sup>-</sup>/Å<sup>2</sup>) | 76.0 | Model validation |  |
| Defocus range (μm) | -1.0 to -2.5 | MolProbity score | 0.89 |
| Micrographs collected | 9,511 | Clash score | 1.51 |
| Pixel size (super resolution) |  | C-beta outliers (%) | 0 |
| initial (Å) | 0.435 | Rotamer outliers (%) | 0 |
| calibrated (Å) | 0.416 | Ramachandran favored (%) | 99.69 |
| Map reconstruction |  |  |  |
| Image processing package | RELION 3.1 | Resolution (Å) |  |
| Extracted particles |  | 0.143 GSFSC unmasked | 3.7 |
| for 2D classification | 996,546 | 0.143 GSFSC spherical mask | 3.6 |
| for 3D classification | 391,668 | 0.143 GSFSC tight mask | 3.5 |
| final count | 236,728 | 3DFSC sphericity (unmasked) | 0.985 |

**Supplementary Figure 1.**
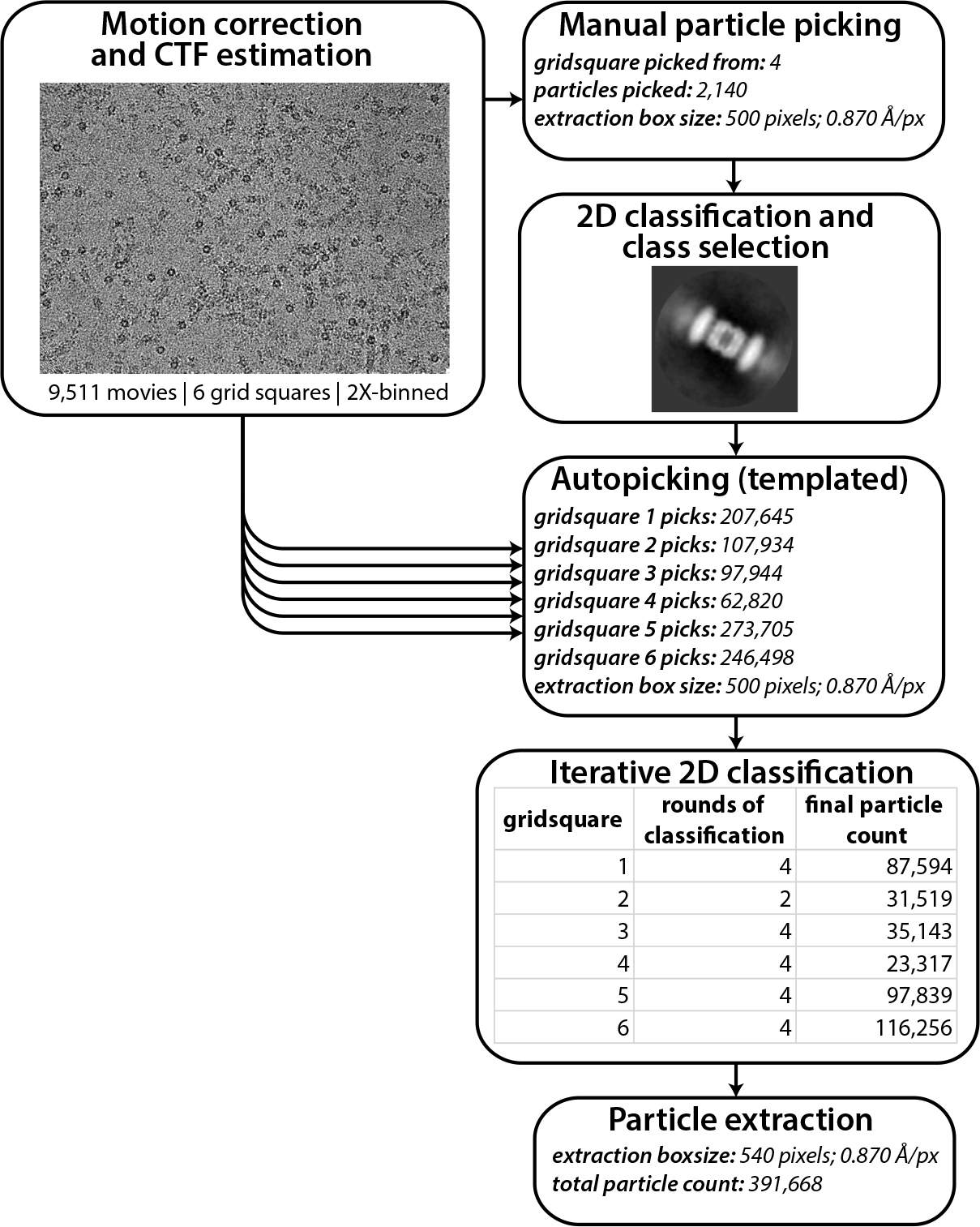
Pre-processing workflow. Initial micrograph and 2D particle processing. Job types and key parameters noted.

**Supplementary Figure 2.**
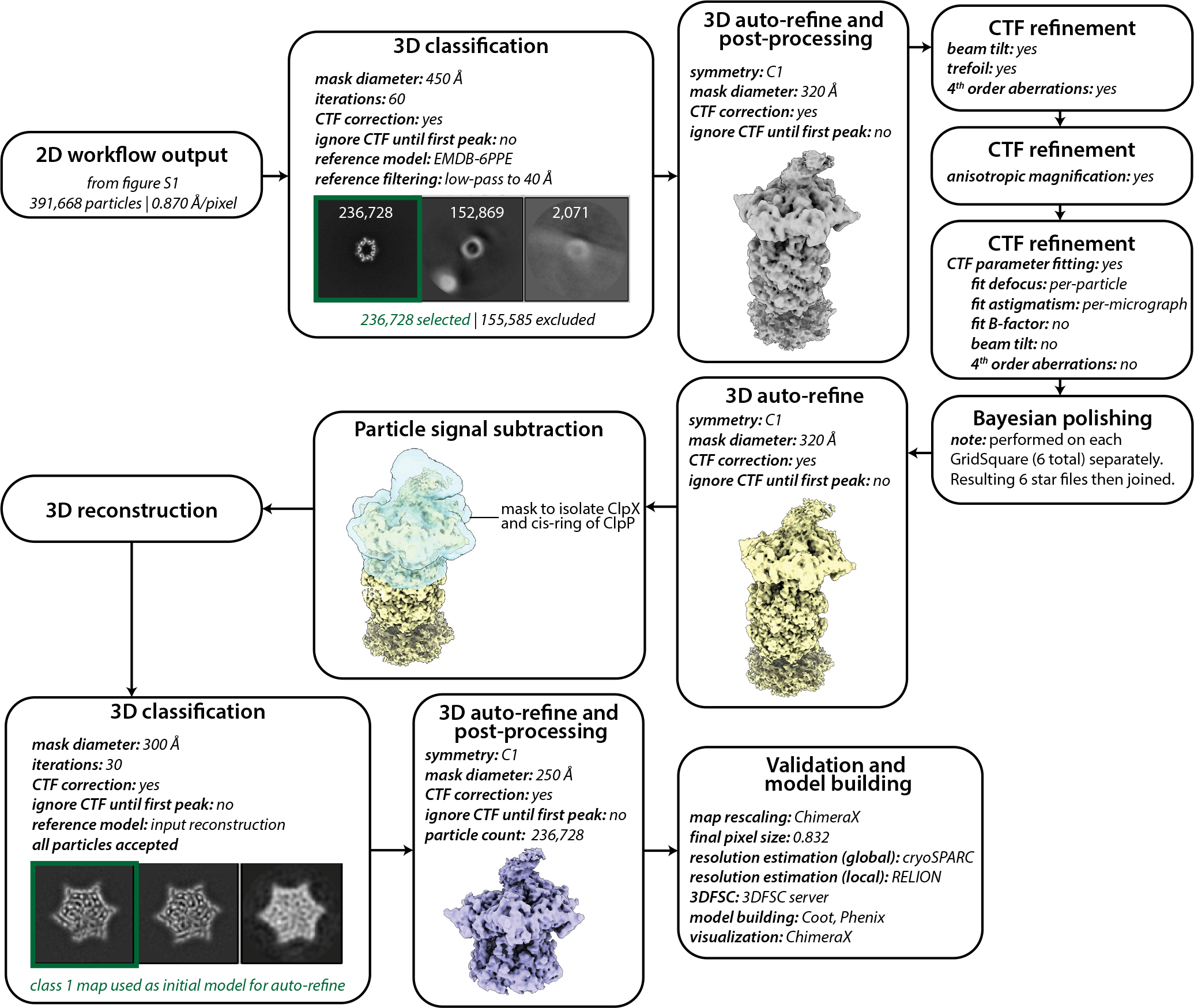
RELION 3D processing workflow. Job names, job details, and non-default parameters are noted in each box.

**Supplementary Figure 3.**
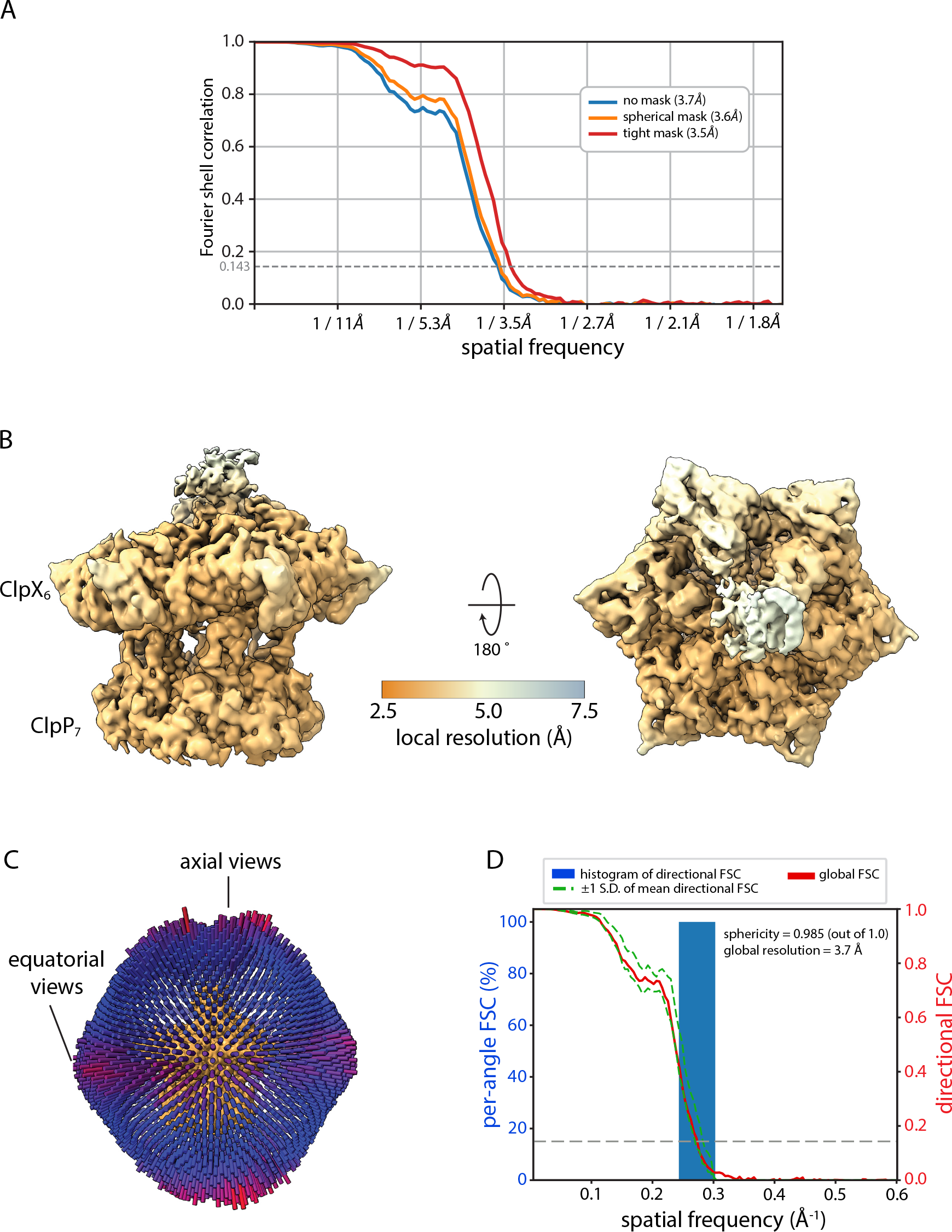
Map assessment and validation. (**A**) Global resolution estimation by GSFSC implemented in CryoSPARC. (**B** Local resolution assessment as estimated by RELION. (**C**) Projection angle distribution and directional FSC as estimated by the 3DFSC server (Tan *et al*. 2017).

